# Mutations in *ErbB2* accumulating in the male germline measured by error-corrected sequencing

**DOI:** 10.1101/2024.08.14.607923

**Authors:** Atena Yasari, Monika Heinzl, Theresa Mair, Tina Karimian, Shehab Moukbel Ali Aldawla, Ingrid Hartl, Andrea J. Betancourt, Peter Lanzerstorfer, Irene Tiemann-Boege

**Affiliations:** Institute of Biophysics, Johannes Kepler University, Linz, Austria; University of Applied Sciences Upper Austria, School of Engineering, Wels, Austria; Department of Evolution, Ecology and Behaviour, University of Liverpool, Liverpool, UK

**Keywords:** Duplex sequencing, ultrarare mutation, de novo mutations, driver mutations, germline mutagenesis, receptor tyrosine kinase, *ErbB2*

## Abstract

Mutations in the male germline are a driving force behind rare genetic diseases. Driver mutations enjoying a selective advantage expand to mutant clusters within the aged testis and are thus overrepresented in sperm with age. Other kinds of driver mutations, occurring prepubescently, have been the focus of recent attention given their high occurrence rate independent of age. Here, we investigated the ErbB2 gene via error-corrected sequencing and detected a high percentage of missense mutations, including recurrent mutations, which were observed mainly in the tyrosine kinase domain with likely functional consequences, as we verified for a subset via biophysical methods. While these mutations increased with age, we found no evidence that they originated from mutational clusters in the aged testis, and young donors also showed an accumulation of driver mutations, suggesting that mutational enrichment is not exclusive to the sexually mature germline but can occur earlier during germline development, and are likely evenly distributed in the testis and stable in size.

## Introduction

Driver or selfish mutations, which are common in cancer, induce higher rates of self-propagation in the cells that carry them, resulting in subclonal expansion events that increase with age (Yates et al. 2015; Jamal-Hanjani et al. 2017; Turajlic et al. 2018; Turajlic et al. 2019); thus, self-propagation can increase within an organism due to positive selection (Yates et al. 2015; Jamal-Hanjani et al. 2017; Loeb et al. 2019; Turajlic et al. 2019), following similar rules as those used for other species (Lewontin 1970). New or de novo mutation (DNM) events are rarer in the germline than in somatic tissue, possibly due to more robust DNA repair mechanisms (Moore et al. 2021). However, once germline DNMs occur, they might also exhibit driver mutations and lead to subclonal expansions, as is well known to occur in the mature male germline for mutations in a handful of genes in the receptor tyrosine kinase (RTK) pathway, such as FGFR2, FGFR3, HRAS, PTPN11, KRAS, RET, BRAF, CBL, MAPK1, MAPK2, and RAF1 (Tiemann-Boege et al. 2002; Qin et al. 2007; Choi et al. 2012; Giannoulatou et al. 2013; Shinde et al. 2013; Yoon et al. 2013; Maher et al. 2014; Maher et al. 2016; Maher et al. 2018; Moura et al. 2024; Striedner et al. 2024). These DNMs (also known as selfish (Goriely and Wilkie 2012), RAMP (Arnheim and Calabrese 2016) or paternal-age effect (PAE) (Crow 2012) mutations are often missense and gain-of-function changes associated with rare congenital disorders.

Until recently, these driver mutations have been hypothesized to expand exclusively in the sexually mature male germline. Specifically, they result in modified functionality of the protein usually associated with hyperactivation of the RTK signaling pathway (Naski et al. 1996; Li and Hristova 2006; He and Hristova 2008; Krejci et al. 2008; Foldynova-Trantirkova et al. 2012; Ornitz and Itoh 2015; Sarabipour and Hristova 2016; Hartl et al. 2023; Moura et al. 2024). This hyperactivation of RTK signaling might confer a selective advantage to spermatogonial stem cells (SrAp) through changes in symmetric cell division patterns (Choi et al. 2008; Choi et al. 2012; Shinde et al. 2013; Yoon et al. 2013; Eboreime et al. 2022; Moura et al. 2024), as reviewed previously (Arnheim and Calabrese 2009; Arnheim and Calabrese 2016). Consequently, driver mutations accumulate with ongoing cell divisions in the mature male germline, form focal mutation pockets in the dissected aged testis (Qin et al. 2007; Choi et al. 2008; Choi et al. 2012; Shinde et al. 2013; Yoon et al. 2013; Maher et al. 2014; Maher et al. 2016; Maher et al. 2018; Eboreime et al. 2022; Moura et al. 2024; Striedner et al. 2024) and are enriched in the sperm of older donors (Tiemann-Boege et al. 2002; Qin et al. 2007; Goriely et al. 2009; Yoon et al. 2009; Choi et al. 2012; Shinde et al. 2013; Yoon et al. 2013; Maher et al. 2018; Salazar et al. 2022; Moura et al. 2024; Striedner et al. 2024). Correspondingly, the risk of a germline-dominant genetic disorder in offspring increases with paternal age (Arnheim and Calabrese 2009; Crow 2012; Goriely and Wilkie 2012; Arnheim and Calabrese 2016).

However, driver mutations might accumulate earlier, before postpubertal spermatogenesis (Moura et al. 2024). This early accumulation is expected to lead to the formation of germline micromosaics, which could result in a high mutational load in sperm that is independent of the age of the donor (Yang et al. 2021; Moura et al. 2024), as was also described for DNMs from the same parent shared among siblings in pedigrees (Rahbari et al. 2016; Gao et al. 2019). Consequently, the risk of recurrence in siblings or the incidence in the general population might be greater for driver mutations that are already present as micromosaics at a young age. This is particularly worrisome since most RTK driver mutations are associated with rare genetic disorders or cancer. Consequently, it is imperative to understand this type of mutagenesis and the expansion patterns of driver mutations in the male germline.

One gene of particular interest is erb-b2 receptor tyrosine kinase (*ErbB2*), a member of the epidermal growth factor receptor family, which also includes *EGFR*, *ERBB3*, and *EBBB4* and regulates RTK signaling. Mutations in *ErbB2* have been detected in numerous somatic tumors of the breast, ovary, lung, large intestine, and prostate (Shin et al. 2011; Subramanian et al. 2019) and have also been implicated in embryonal carcinoma of the testes and advanced testicular teratomas (Shin et al. 2011). In the mature testis, *ErbB2* is broadly expressed in spermatogonia, early spermatocytes, elongating/elongated spermatids, Sertoli cells, and Leydig cells (Guo et al. 2018; Guo et al. 2020) and potentially regulates signaling during mitosis and the onset of meiosis in germ cells and spermiogenesis (Shin et al. 2011). Like in other RTK mutations, missense mutations in *ErbB2* might have a selective advantage and expand in the male germline.

Here, we investigated the degree to which ErbB2 accumulates in the sperm DNA of differently aged donors via error-corrected sequencing (ecSeq), also known as duplex sequencing (DS). This approach allowed us to collect a large number of DNMs, compare substitution frequencies and mutation types in the coding regions of *ErbB2,* and detect instances of positive selection. We also investigated the expansion patterns of aged dissected testes and the effect of selected mutations on receptor signaling to better understand the functional changes associated with these substitutions.

## Results

### Targeted error-corrected sequencing to detect mutations

DS or ecSeq has the lowest reported error rate for Illumina sequencing, achieved by using a double-barcode strategy that assembles information from both strands of the original DNA molecule (Schmitt et al. 2012; Kennedy et al. 2014) and reviewed previously (Salk et al. 2018). Consequently, DNA lesions or PCR errors can be distinguished from true mutations (Arbeithuber et al. 2016); thus, this method is considered one of the advanced approaches for detecting low-frequency mutations. However, nicks in DNA resulting from random DNA shearing accompanied by high-energy bursts of sonication combined with error-prone repair of the templates before adaptor ligation might still be sources of artifacts. Specifically, the removal of nicks by strand extension and repair of protruding 3’DNA ends during library preparation eliminate the original duplex sequence information necessary to distinguish real mutations from DNA lesions (Abascal et al. 2021).

For this reason, we implemented experimental ‘repair steps’ during library preparation. Specifically, we used dideoxy bases (ddBTPs) that block nick extensions and Mung Bean blunt endings to reduce errors derived from DNA synthesis by the enzyme mixture used before adapter ligation, as described previously (Abascal et al. 2021). In addition, we treated the DNA with enzymes (Fpg and UNG) to remove the most common DNA lesions (oxo-G and nonmethylated cytosine deamination, respectively). Figure 1 shows a lower average substitution frequency for the 17 repaired libraries (Supplementary Table S1) than for the 25 unrepaired ones (Supplementary Table S2), with comparable substitution frequencies among individual libraries (Supplementary Figure S1). Five of the donors were the same for the repaired and unrepaired libraries, and the others were matched in terms of age; however, the donors differed in terms of sperm quality (50% vs 100% normospermic donors, respectively).

**Figure 1.**
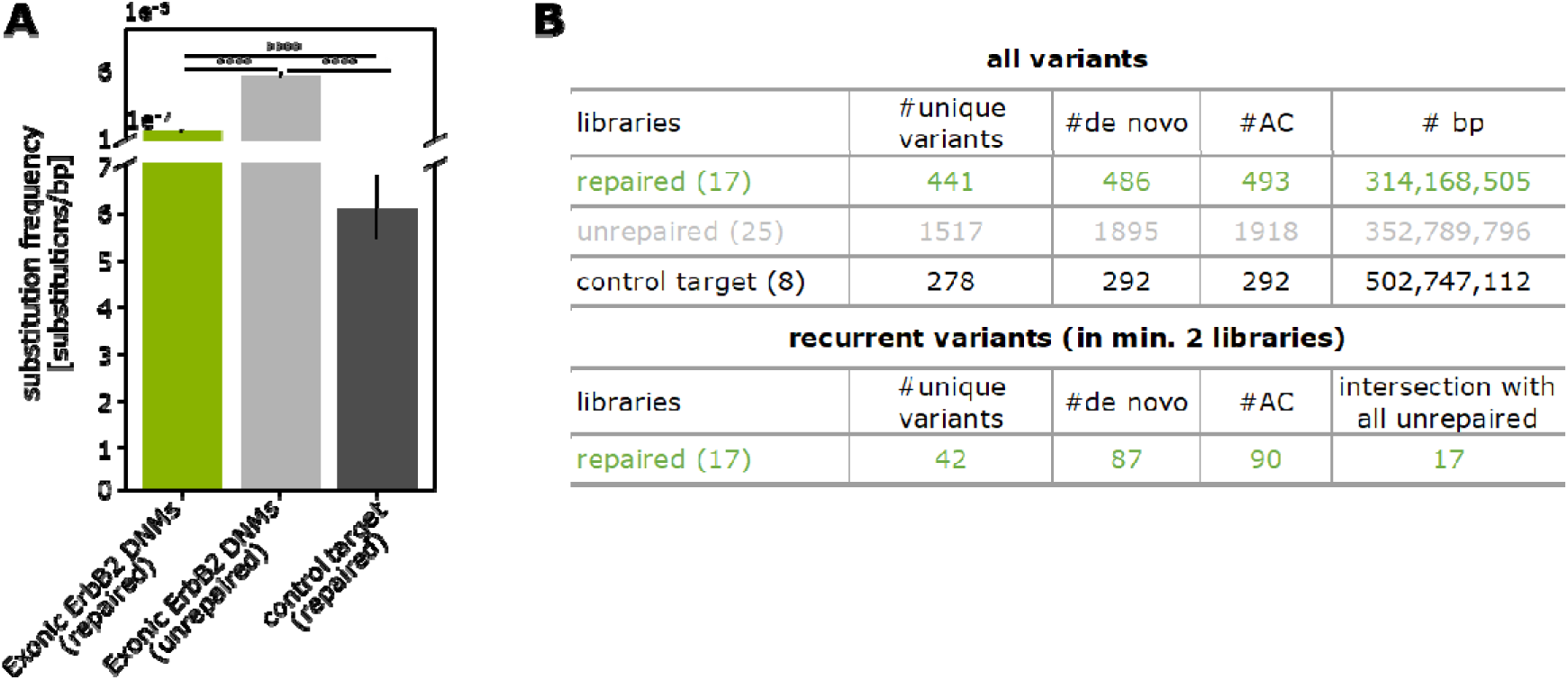
**(A)** Overall substitution frequencies observed in repaired libraries (17 libraries), unrepaired libraries (25 libraries), and negative controls (8 libraries). The high-quality variants (tier 1.1-2.5) listed in Supplementary Tables S1-4 were included. For these estimates, we considered de novo variant counts defined as the sum of all variants observed in the different donors but counted only once if they occurred within the same donor. Note that each of the repaired libraries represents a different donor. For pairwise testing, we used the chi-square test with Bonferroni–Holm correction, and only significant differences are shown. (∗) P value < 0.05; (∗∗∗∗) P value < 0.0001. **(B)** Number of substitutions captured in the different library protocols classified based on the substitution type (unique), the substitution counted in different donors (de novo), or all substitutions occurring within the same or different donors (allele count-AC). For the repaired and unrepaired libraries, we excluded intronic variants. We excluded SNPs (VAF ∼50%) from all the libraries, variants occurring in more than 40% of the libraries or rare haplotypes. The number of sequenced base pairs was estimated by the average coverage in exonic regions in repaired/unrepaired libraries or targeted regions for the negative controls. ‘Recurrent’ represents variants observed in at least two donors. Of the 42 unique variants found to recur in the repaired libraries, 17 were also observed in the unrepaired libraries (intersection).

For the repaired libraries, we observed a reduction in the frequency of C>A/G>T transversions, a common substitution for oxo-G lesions (Supplementary Figure S2). The cosine similarity values comparing the mutational spectra with exonic variants in COSMIC or gnomAD were also more similar for repaired libraries than for unrepaired libraries (0.85 vs 0.71-COSMIC or 0.79 vs 0.64-gnomAD, respectively). We suspect that unrepaired libraries had more artifacts that were reduced in the ‘repaired’ libraries by the nick- and end treatments and the enzymatic removal of lesions. We also used a lower sonication energy (a potential source of nicks) for the repaired libraries. Based on these results, data from unrepaired libraries were not used further except if specified in individual analyses.

To further validate the ecSeq methodology, we measured the overall substitution frequency of a control human genomic region in 8 different donors (7 of these donors were also analyzed for exonic *ErbB2* regions), and the data are shown in Supplementary Table S3. For these controls, we also used the “repaired library preparation” strategy. The control region was a subset (∼10 kb; similar in size to the *ErbB2* region) of a mutagenic test used in other projects (Valentine et al. 2020; Wang et al. 2021) that comprises different genomic sequences, each 0.5 kb in size (total 20 kb), with no known evidence of positive/negative selection or a functional role; additionally, the control region was designed to represent a balanced sequence context in terms of GC content and genic/nongenic, coding/noncoding regions.

Using the ‘repaired’ approach, we measured a substitution frequency of 5.8×10^-7^ in these controls (Figure 1A). A comparable substitution frequency of 1.9-5.5×10^-7^ was reported for a human airway cell line (Wang et al. 2021) and a human lymphoblastoid cell line (Cho et al. 2023) targeting the 20 kb control targets.

The threefold lower substitution frequency measured in the negative controls compared to the *ErbB2* region (5.8×10^-7^ vs. 1.5×10^-6^, respectively) suggested that two-thirds of the DNMs were exclusive to a process within *ErbB2*, but we cannot unambiguously exclude a contribution from artifacts or contaminating somatic cells. Contaminating somatic cells, in particular, may increase the substitution frequency, given that they have 10- to 100-fold greater mutation rates than germline cells (Abascal et al. 2021; Moore et al. 2021). In normospermic individuals, the percentage of nonsperm cells in semen typically ranges from 1-10%, but nonsperm cells vary with factors such as fertility, age, health, and lifestyle and are potentially more prevalent in cases of infertility (Bjorndahl et al. 2003; Carlsen et al. 2004). Among the sperm donors examined for both exonic *ErbB2* targets and controls, the difference in substitution frequency between *ErbB2* and control regions was also threefold (Supplementary Figure 1D).

We also implemented a series of filtering steps at the bioinformatic level. These included the assignment of a tier classification (tiers 1.1 to 7) by the Variant Analyzer (VAR-A), which verified the reliability of the variant call based on evidence such as mate information, family size, and errors within a family (Povysil et al. 2021). We further carried out a haplotype analysis and identified variants that cooccurred with another rare variant or rare haplotype (Supplementary Figure S3). These rare haplotypes were identified in multiple libraries (both within and among repaired libraries), were usually short, had ≤ 10 base pairs, and were sometimes tagged by supplementary alignments (part of the read mapped to a different region of the genome). Since it is unclear whether these variants were exclusive to *ErbB2*, we removed these variants from further analyses. We also removed variants within a short tandem repeat (STR), given the problematic and correct variant annotations at these sites.

In total, we measured 493 substitutions (high-quality tier of 1.1 to 2.5) in the exonic region of *ErbB2* (4557 bp) in the repaired libraries (Figure 1B). Of those, 486 substitutions were classified as de novo occurring in different donors (in at least two libraries), and 441 were unique substitutions. Those variants observed in multiple donors were labeled recurrent (Figure 1; Supplementary Table S4). Of the 42 recurrent variants in the repaired libraries, 2 variants cooccurred in more than 2 donors, and 40 variants occurred in two different donors. Of the recurrent variants, 17 were also observed in the unrepaired libraries (Figure 1B). No significant differences in age, age, diagnosis, absolute number of sperm, volume (ml), or MS/ml were detected between the recurrent and nonrecurrent variants based on the Mann⍰Whitney U test (p value< 0.05; Supplementary Table S5-S6).

### A greater number of substitutions was observed in older donors

We analyzed libraries from three age classes: six from younger donors (19 to 30 years old), six from middle-aged donors (31 to 45 years old), and five from older donors (46 to 63 years old); all healthy individuals, mainly from individuals of European ancestry. The average coverage depth of the repaired libraries was ∼5000x (max ∼7000x). We observed that the substitution frequency increased with age (Figure 2A) and was significantly greater in the middle-aged and older groups than in the younger group. This difference between age groups was not as pronounced with respect to the recurrent dataset, probably because of the reduced power of a smaller sample size.

**Figure 2:**
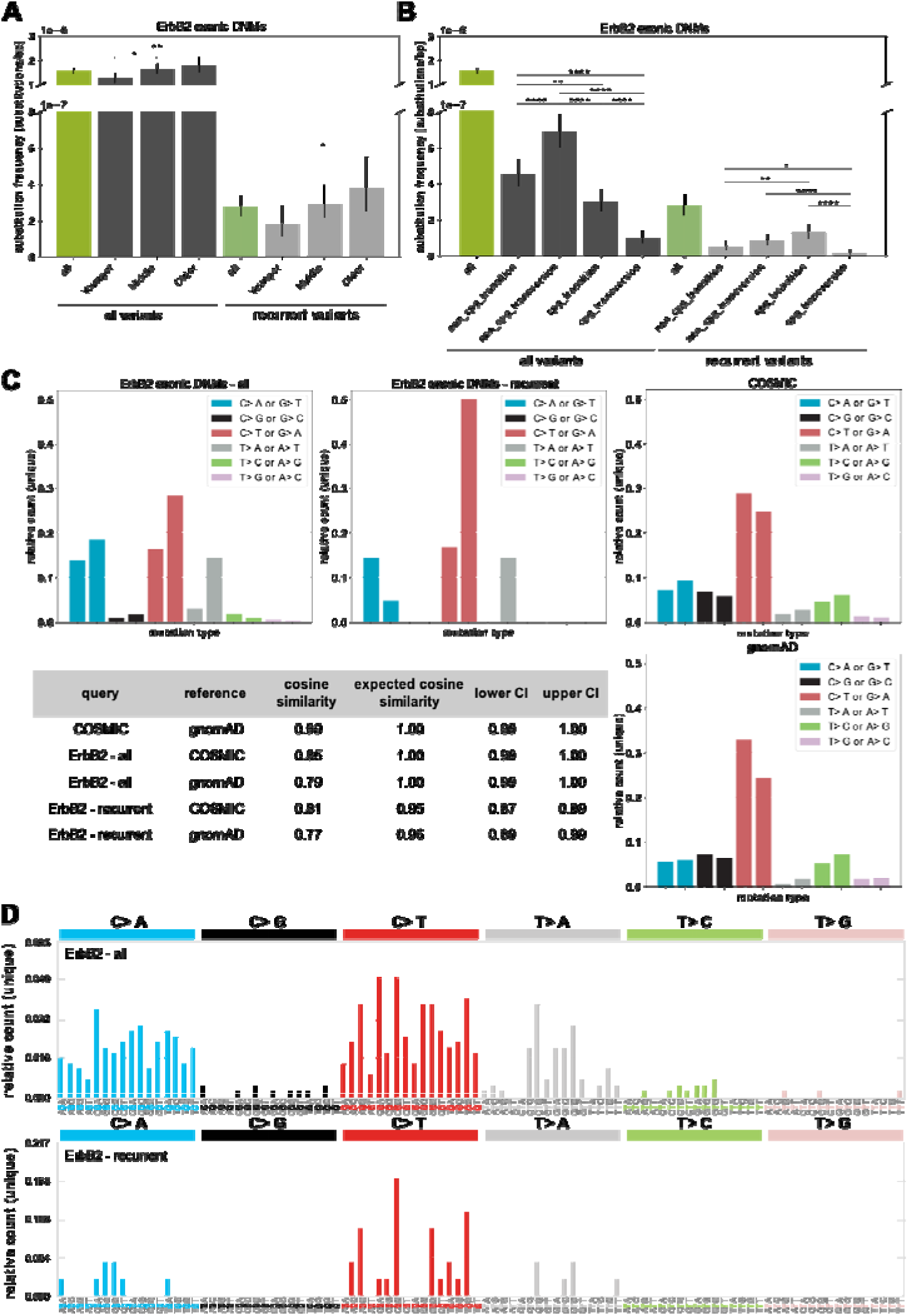
Analysis of substitutions in *ErbB2*. **(A)** Substitution frequencies of all- (n=486) and recurrent (n=87)-exonic variants identified in different donors (de novo) classified into different age categories (younger, middle-aged, older) in the repaired libraries. **(B)** Substitution frequency of all- and recurrent-exonic variants classified into substitution types. The high-quality variants (tier 1.1-2.5) listed in Supplementary Tables S1 and S4 were included. **(C)** Mutational substitution spectra categorized after the untranscribed strand. **(D)** Signature within the context of the two flanking bases of all- (n=441) and recurrent-unique (n=42) variants of the repaired libraries compared to the spectra extracted from COSMIC and gnomAD. For pairwise testing, the chi-square test with Bonferroni–Holm correction was used, and only significant differences are shown in A and B. (∗) P value < 0.05; (∗∗∗∗) P value < 0.0001

We further analyzed the mutational spectra, mutational signatures, and transcriptional bias of the observed substitutions. When considering all the data, the most frequent substitution types were non-CpG transversions and non-CpG transitions (Figure 2B). The mutational spectra also reflected a high number of G>T/C>A or (S>W) transversions and G>A/C>T or (S>W) transitions. Based on the cosine similarity value, the substitutions in *ErbB2* were more similar to the variants reported in tumors for *ErbB2* (COSMIC) than to the variants identified in the general population (gnomAD), as shown in Figure 2C (0.85 vs. 0.79, respectively). The mutational signature comparing the variants also in the context of the 5⍰ and 3⍰ adjacent bases indicated that most substitutions occurred in the context of an S-C-R trinucleotide, and this signature overlapped by ∼52% with SBS4 (associated with tobacco smoking), ∼22% with SBS30 (deficiency in base excision repair due to inactivating mutations in NTHL1), ∼15% with the SBS1 pattern (spontaneous or enzymatic deamination) and ∼11% with SBS5 (unknown etiology; Supplementary Figure S4). We have no knowledge of whether the donors had a particular disease background or smoking history.

In contrast, when considering only recurrent variants, we observed mainly S>W transitions, mostly at CpG sites (Figure 2B), consistent with a higher percentage of 5meC>T mutations. Additionally, for this dataset, the mutational spectra of recurrent variants were also more similar to those of COSMIC than to those of gnomAD (0.81 vs. 0.77, respectively). Curiously, the mutational spectra of recurrent variants were also similar to those of another driver gene (*FGFR3*) characterized by ecSeq in the male germline (Salazar et al. 2022) (cosine similarity between *ErbB2* and *FGFR3* of 0.89, CI = 0.89-0.99).

The mutational signature for recurrent variants showed the most frequent substitutions in the context of V-C-G sites, with ∼26% overlap of variants explained by the SBS1 pattern (spontaneous deamination of 5-methylcytosine), ∼28% by SBS5, and ∼46% by SBS87 (thiopurine chemotherapy treatment; Supplementary Figure S5). When analyzing which substitutions accumulated more often in the genic region, we found that strong to weak transversions (C>A) and transitions (C>T) and pyrimidine to purine transversions (T>A) occurred more often in the transcribed strand than in the untranscribed strand (Supplementary Figure S6).

### Enrichment of recurrent substitutions in the extracellular and protein kinase domains

Driver mutations in the male germline might also be associated with a functional change or a change in the signaling activity of the receptor (Arnheim and Calabrese 2009; Goriely and Wilkie 2012; Arnheim and Calabrese 2016). To explore this possibility, we examined where the substitutions occurred within the ErbB2 protein domains. Figure 3A shows that when considering all the data, most of the substitutions occurred in the transmembrane domain and flanking regions, with one-third of the substitutions predicted to result in a deleterious change based on the SIFT scores (Ng and Henikoff 2003; Sim et al. 2012); this change is one of the most commonly used algorithms for predicting the effect of a substitution on protein function (Supplementary Table S1). A small fraction of the DEGs were associated with cancer (according to the COSMIC database).

**Figure 3:**
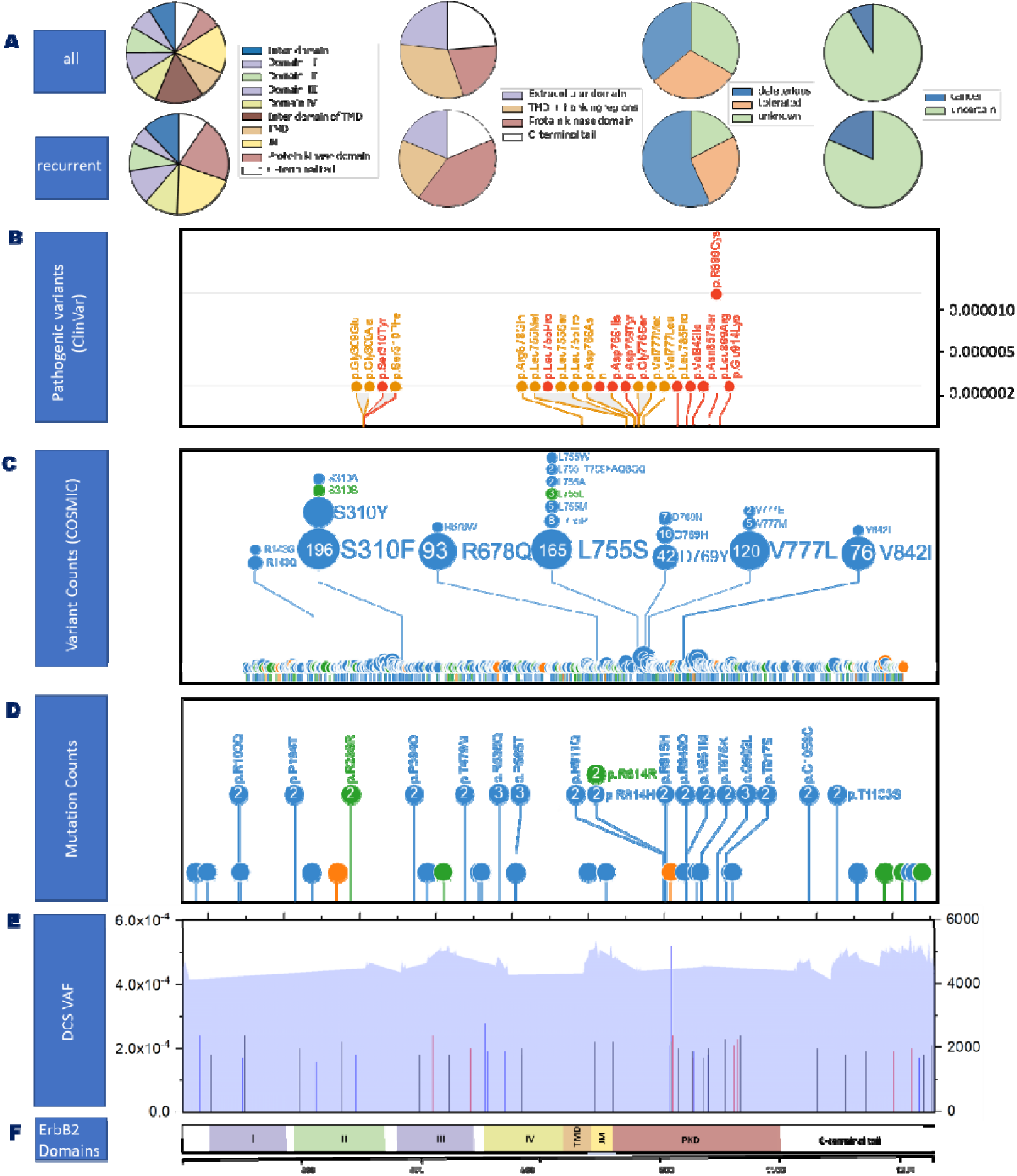
Distribution of substitutions in different protein domains of ErbB2. **(A)** Substitution frequencies split by domain and deleteriousness based on the SIFT score (Sim et al. 2012) and correlation with cancer based on COSMIC of de novo variants found in the repaired (486) and recurrently repaired (87) datasets detected in all 17 repaired libraries (variants listed in Tables S1 and S4). **(B-D)** Distribution of pathogenic/likely pathogenic missense substitutions in *ErbB2* correlated with germline or somatic cancers reported in ClinVar with pathogenic variants in red and likely pathogenic variants in orange paired with reference data from the Exome Aggregation Consortium (ExAC) of gnomAD v3.1 access Jan2024 (Karczewski et al. 2020) displayed on the right y-axis **(B)**, the number of reported exonic somatic amino acid substitutions in *ErbB2* associated with tumors retrieved from the COSMIC v96 database (Liu et al. 2019) **(C)**, and the recurrent exonic mutations (42) with their respective VAF measured in 17 different sperm DNA libraries with numbers displayed representing randomly selected donors with the recurrent variant **(D)**. The plots were prepared with protein paint (Zhou et al. 2016), with the size of the ball proportional to the number of mutation hits/counts and the color in C-D denoting the substitution type: missense, nonsense, or silent mutation (blue, orange, green, respectively). **(E)** Lines represent the variant allele frequency (VAF) of the 42 recurrent exonic *ErbB2* variants scaled to the left y-axis. Note that variants occurring in multiple donors were plotted using the mean VAF. Variants also reported in COSMIC are shown as red lines, and those in gnomAD are shown as blue lines. The gray area represents the median DCS coverage, with the DCS counts displayed on the right y-axis. **(F)** ErbB2 protein domains. (I) Domain I, (II) domain I, (III) domain III, (IV) domain IV, (TMD) transmembrane domain, JM, and protein kinase domain of *ErbB2*.

Variants that occurred more than once independently (i.e., in different donors) appear to be more likely to be responsible for clonal expansion than all the exonic *ErbB2* DNMs. First, most (34 out of 42) of the recurrent variants were missense substitutions, 6 were synonymous, and 2 were classified as stop-gain mutations (Table 1). Second, many of these recurrent mutations are likely to be deleterious. Approximately one-fifth of these recurrent substitutions were reported as somatic cancer-promoting mutations in COSMIC (compared to 8% considering all variants), and half were reported in gnomAD and are likely viable; 10 have never been reported before. Furthermore, ∼50% of these recurrent substitutions are highly likely to be deleterious according to their CADD score (Kircher et al. 2014) (Table 1 and Supplementary Table S4). Third, when classifying the substitutions per domain, two-thirds of the substitutions occurred in the protein kinase domain (PKD) (Figure 3A).

**Table 1.**
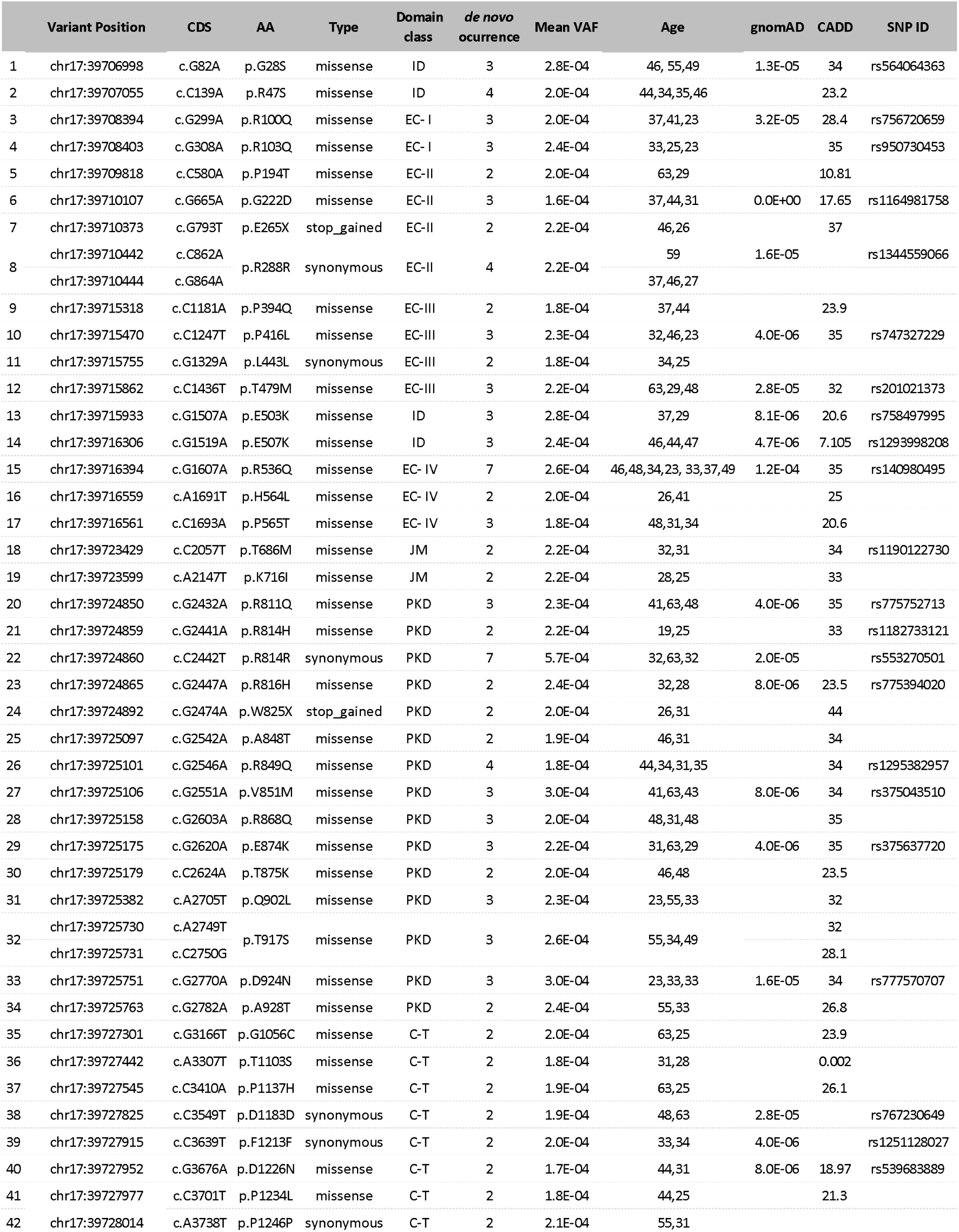
Recurrent variants found in young, middle-aged or older donors are displayed together with the respective mean VAF of only repaired or both repaired and unrepaired libraries; if present, the number of total observations in different donors (both repaired and unrepaired libraries) are classified into interdomain (ID), extracellular domain I-IV (EC I-IV), juxtamembrane domain (JM), protein kinase domain (PKD), and C-terminal tail (C-T).

Variants categorized as associated with a disorder by ClinVar or reported in COSMIC also mainly accumulated in the PKD gene of ErbB2 (Figure 3B-C), as was also observed for recurrent DNMs (Figure 3D), for which the mutation frequency or variant allele frequency (VAF) was ∼2×10^-4^ (Figure 3E); similarly, the highest average VAF (5.2×10^-4^) was measured for c.C2442T (p.R814R), a silent mutation in the kinase domain (see Table 1 and Supplementary Table S4). This enrichment in the kinase domain of pathogenic and tumor-related variants and recurrent DNMs in sperm suggest that this domain is a target for mutations likely associated with clonal expansion in ErbB2.

### Evidence for positive selection: passenger versus driver mutations

To investigate whether the variants enriched in sperm are driven by positive selection, we performed a version of a *d*_N_/*d*_S_ analysis, which compares nonsynonymous vs. synonymous substitutions adjusting for the local sequence context (Martincorena et al. 2017). A *d*_N_/*d*_S_ ratio close to the reference value of one (1) indicates no selection, that below the reference value implies negative selection, and that above the reference indicates positive selection (Nielsen 2005; Martincorena et al. 2017). Table 2 shows that *ErbB2* substitutions have a signature of positive selection with a *d*_N_/*d*_S_ ratio of 1.5, which is significantly greater than one. Curiously, this ratio increased considerably when considering only recurrent substitutions (*d*_N_/*d*_S_ = 3.7), further supporting that recurrent mutations are a subset of variants enriched for targets of positive selection. For both datasets, the strongest indication for positive selection was in the protein kinase domain (Table 2).

**Table 2.** Estimates of the *d*_N_/*d*_S_ ratio following (Martincorena et al. 2017) calculated in total or per protein domain (EC: extracellular domain, TMD + flanking regions, PKD: protein kinase domain, C-terminal tail) and per age group for de novo exonic *ErbB2* variants (n=486 mutations for the repaired libraries, n=457 nonsynonymous and synonymous variants, or n= 87 recurrent mutations; n=83 nonsynonymous and synonymous). Variants were retrieved from Supplementary Table S1 and S4, omitting variants resulting in stop codons. The age category was defined as sperm donors younger than 30 years, middle-aged donors between 30 and 45 years and older donors older than 45 years. This partitioning ensured similar sample sizes between groups.

|  |  | de novo |  |  |  | de novo recurrent |  |  |  |  |
| --- | --- | --- | --- | --- | --- | --- | --- | --- | --- | --- |
|  |  | *n*<br>*nonsyn* | *n* *synon* | *dN/dS* | *p*<br>*value* |  | *n*<br>*nonsyn* | *n* *synon* | *dN/dS* | *p*<br>*value* |
| All |  | 326 | 131 | 1.5 | 2E-10 | * | 71 | 12 | 3.7 | 0E+00 * |
| Domain | EC | 163 | 65 | 1.4 | 2E-05 | * | 30 | 4 | 4.6 | 2E-11 * |
|  | TMD | 26 | 14 | 1.1 | 5E-01 |  | 4 | - | - | - |
|  | PKD | 68 | 19 | 2.1 | 4E-08 | * | 27 | 2 | 7.5 | 3E-15 * |
|  | C-terminal | 69 | 33 | 1.3 | 7E-02 |  | 10 | 6 | 1.2 | 6E-01 |
| Age Class | Younger | 85 | 38 | 1.3 | 3E-02 | * | 16 | 1 | 8.5 | 2E-10 * |
|  | Middle | 156 | 60 | 1.5 | 7E-06 | * | 34 | 6 | 3.9 | 1E-10 * |
|  | Older | 85 | 33 | 1.6 | 3E-05 | * | 21 | 5 | 2.5 | 3E-04 * |

Mutations in all age groups also showed signatures of positive selection, with similar values observed for the younger, middle and older groups; these values slightly increased with age when considering all the data (Table 2). Given the smaller sample size of recurrent variants, the *d*_N_/*d*_S_ estimates for these mutations have large confidence intervals, but all categories analyzed show statistical evidence for positive selection.

### Spatial distribution of selected variants in the male gonad

To gain further insight into the accumulation of *ErbB2* mutations in the germline of sexually mature males, we screened the spatial distribution of three variants in the postmortem testes of two different donors, 70-year-old and 73-year-old individuals. In particular, we examined c.428G>A (p.R143Q), c.2033G>A (p.R678Q), and c.2524G>A (p.V842I), which were selected based on the association of the variant with a cancer phenotype, different clinical significances (pathogenic to likely benign), high predicted deleteriousness (CADD score) (Kircher et al. 2014), COSMIC reports, and differences in embryonic viability of these variants based on gnomAD reports (Figure 4A). Furthermore, p.R678Q is the fourth most common mutation in *ErbB2* reported in COSMIC, and p.V8421 is an activating mutation (Bose et al. 2013). Furthermore, based on our ecSeq data, these variants were found to occur in sperm at VAFs ∼10^-4^ from at least 3 different donors when unrepaired libraries were also considered (Figure 4B). Note that we also observed different nucleotide substitutions at the same codons, rendering alternative missense or silent substitutions, as reported in Supplementary Table S1 and S2.

**Figure 4:**
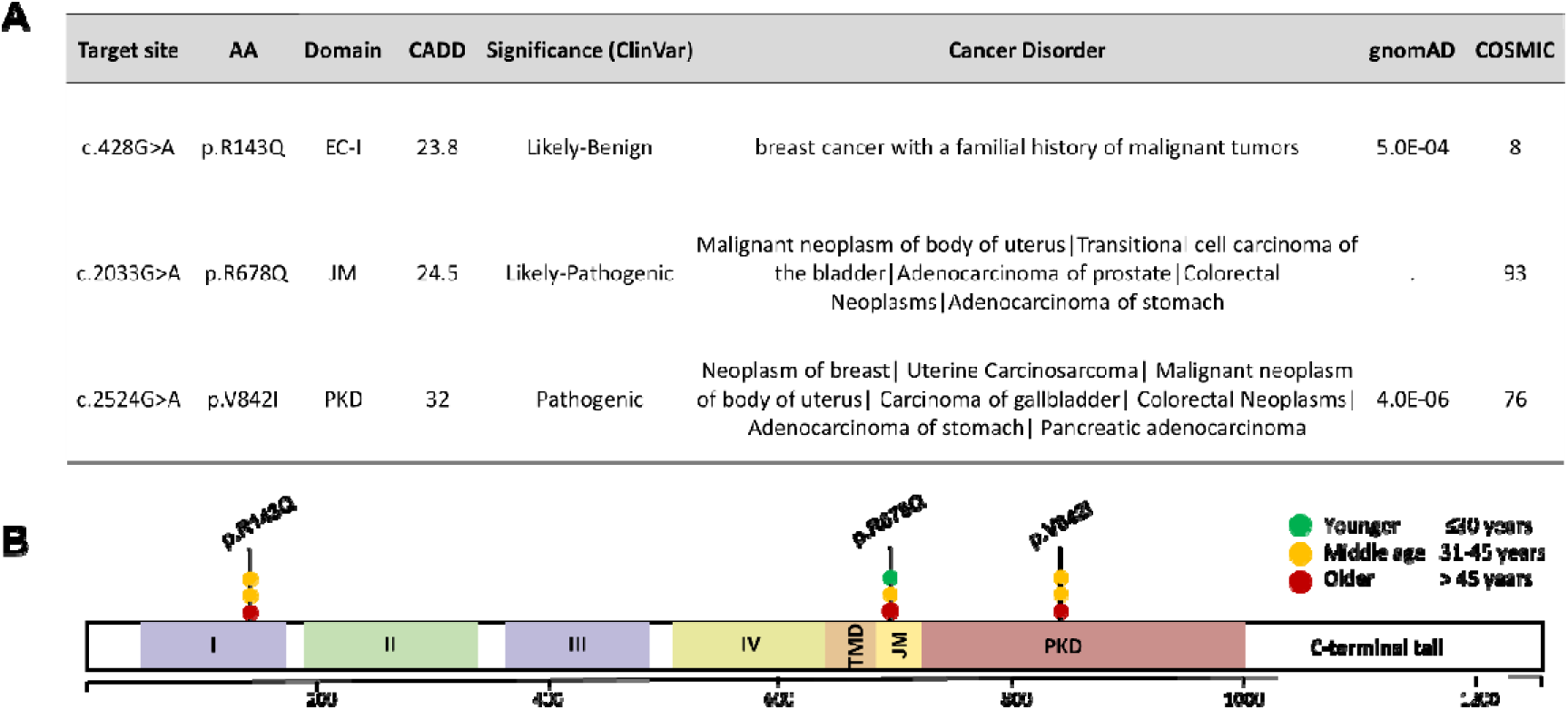
**A.** *ErbB2* variants explored for testis dissection analysis. The target site of the coding sequence position is listed, AA: amino acid. CADD score (Kircher et al. 2014) GRCh38-v1.6 and deleteriousness. VAF: variant allele frequency. All gnomAD information was retrieved from v3.1. (Karczewski et al. 2020). COSMIC data were based on version 94 (Liu et al. 2019). The data were called based on the transcript ENST00000269571 **B.** Relative position of the three selected variants within the *ErbB2* domains and the recurrence of variants in both unrepaired and repaired libraries are shown in Supplementary Table S1 & S2. Each dot represents one mutation count that is color-coded based on the age of the sperm donor. (I) Domain I, (II) domain I, (III) domain III, (IV) domain IV, (TMD) transmembrane domain, juxtamembrane domain (JM), and protein kinase domain (PKD) of ErbB2.

We followed the testis microdissection technique (Qin et al. 2007; Choi et al. 2008; Shinde et al. 2013; Yoon et al. 2013) in combination with digital droplet PCR (ddPCR), as described previously (Moura et al. 2024). Briefly, each testis was divided into 6 slices, and each slice was further partitioned into 32 pieces. The extracted DNA from four adjacent testis pieces was pooled, resulting in 8 pools per slice or 48 pools per testis, as shown in Figure 5A (for details, see (Moura et al. 2024)). For each pool, we screened 270,000-300,000 genomes. With this input, some samples yielded 1-3 mutants, implying VAFs of ∼10^-5^ (Supplementary Table S7). In 15-30% of the samples, we did not detect any mutations (Supplementary Figure S7). Note that at an input level of 300,000 genomes, the chance of finding zero mutations is 60% (based on a Poisson distribution with λ = 0.5); thus, failure to observe positive counts at this depth can still be consistent with mutations present at VAFs lower than the detection threshold.

**Figure 5:**
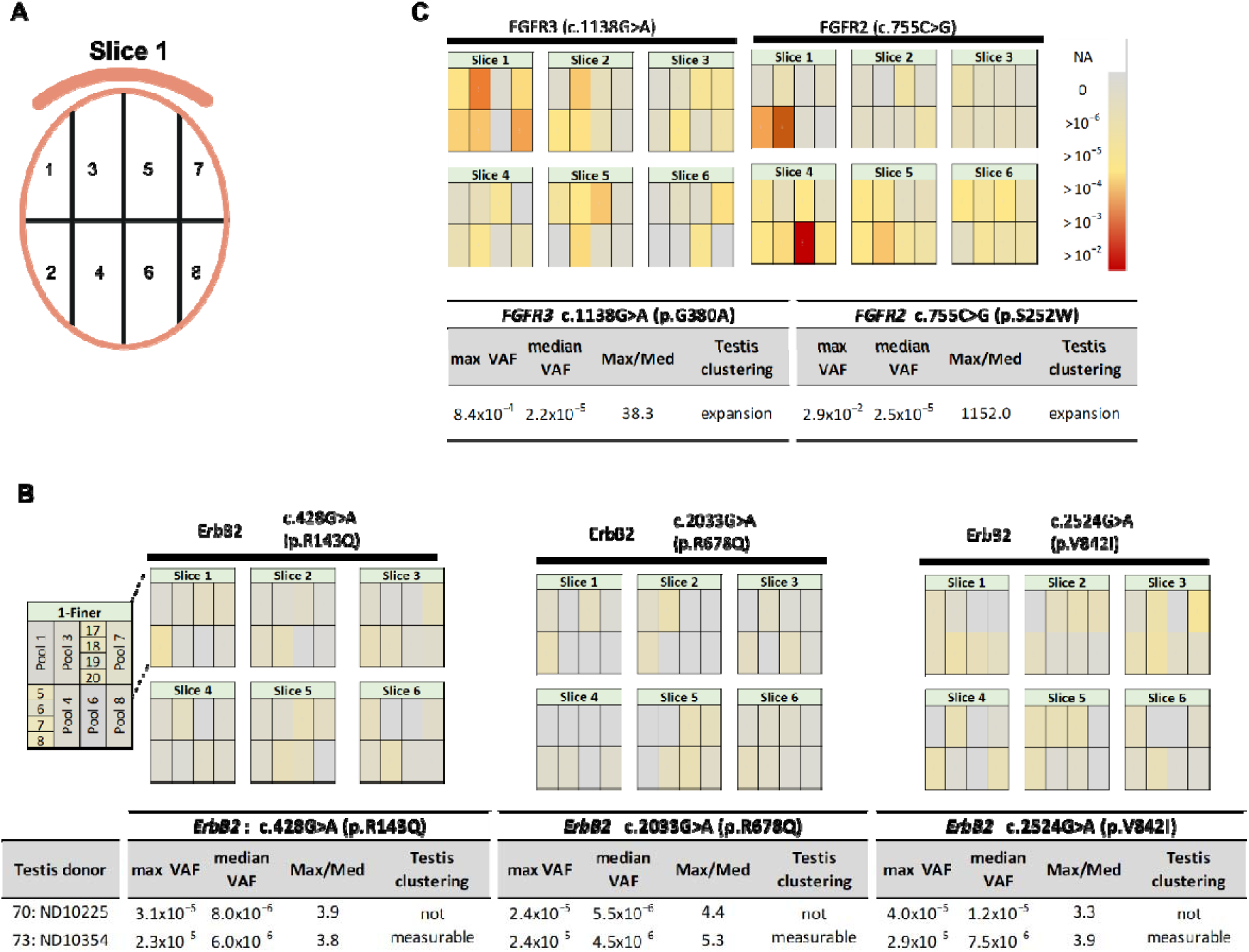
Mutational screening of testis DNA via droplet digital PCR (ddPCR). **A.** Testis cutting scheme strategy: The testis was cut in half and further divided into 6 slices. Each slice was dissected into 32 pieces, and the DNA from the four adjacent pieces was pooled and analyzed via ddPCR (8 pools per slice). The curved line above the slice denotes the epididymis for orientation purposes. **B.** VAFs of the three *ErbB2* variants measured in the postmortem testis of a 70-year-old Caucasian donor (ID: ND10225). The VAF for a second donor (ND10354) is shown in Supplementary Figure S9. Descriptive statistics of the maximum and median VAF in the 48 pools (data in Supplementary Table S7). The ratio Max/Med represents the ‘hotness’ of the mutation cluster and compares the highest VAF (maxVAF) to the remaining pools (median VAF). **C.** Data on canonical driver mutations expanding in the aging male gonad have been documented for *FGFR2* and *FGFR3*: variant c.755C>G was screened in the testis 374-2 of an unaffected 62-year-old donor (Qin et al. 2007), and c.1138G>A was screened in an unaffected 80-year-old donor (ID 57650) (Shinde et al. 2013). The color scale of the VAF magnitude is representative of all the panels.

We validated the ddPCR measurements by screening the flushed epididymis (mostly the sperm of the testis donor) and a piece of scrotal skin, both of which are expected to have different mutation frequencies than individual testis pieces. In addition, we screened the same sperm donors as those used for ecSeq (Supplementary Figure S8; Table S8) to compare the measurements between the two methodologies. For all three variants, the skin measurements (proxy of a negative control) revealed significantly fewer VAFs (except for p.V821I) than did the epididymis (proxy of a positive control). Furthermore, we found good congruence between the VAFs of donor-matched sperm data measured with ecSeq and ddPCR. This finding suggested that the ddPCR method is appropriate for screening expansion clusters if the subclonal expansions observed in *FGFR3* (Shinde et al. 2013) and *FGFR2* (Qin et al. 2007) reach VAFs as high as 8×10^-4^ and 2.9×10^-2^, respectively.

None of the three *ErbB2* sites reached VAFs larger than ∼4×10^-5^ (maxVAF; Figure 5B and Supplementary Figure S9). It is possible that the clusters were diluted by the pooling strategy, but the analysis of individual testis pieces within two selected pools showed that this was unlikely (Figure 5B and Supplementary Table S7). The ‘hotness’ of the focal mutation pockets was assessed by the Max/Med ratio, which compares the highest VAF (maxVAF) to the remaining pools (median VAF) and ranges from ∼3- to 4-fold for both testes and the three different variants. For comparative purposes, we included data on two mutations known to form subclonal clusters in the aging testis that were also collected with the same dissection scheme. Specifically, we selected c.1138G>A (p.G380A) and c.755C>G (p.S252W) of the *FGFR3* and *FGFR2* genes, respectively, which are associated with achondroplasia (Shinde et al. 2013) and Apert syndrome (Choi et al. 2008). Figure 5C shows that these two variants form distinct clusters in the testis, with VAFs one to three orders of magnitude greater than the median VAF measured in the remaining testis (Max/Med). In conclusion, we did not observe large differences in VAFs among pools for *ErbB2* variants in any of the two testes (Figure 5B and Supplementary Figure S9) and have no support that the selected *ErbB2* variants accumulate in the sexually mature gonad with age.

### Analysis of signaling activity of selected ErbB2 variants via biophysical methods

#### Some kinase domain variants increase the recruitment of downstream adaptor proteins

We also investigated the changes in the activation of the three focal *ErbB2* protein variants (p.R143Q, p.R678Q, and p.V842I) at the cellular level. For this purpose, we investigated the recruitment of downstream adaptor proteins using total internal reflection fluorescence (TIRF) microscopy, which analyses receptor⍰adaptor interactions at the cell membrane of live cells (Schwarzenbacher et al. 2008), as has been previously done for FGFR3 (Hartl et al. 2023; Moura et al. 2024) and EGFR (Lanzerstorfer et al. 2014). Cells coexpressing both a cytosolic downstream adapter protein (Grb2 or Shc1 fused to the monomeric red fluorescent protein mRFP) and one ErbB2 variant (tagged with the monomeric green fluorescent protein mGFP) were seeded on micrometer-scale antibody-patterned surfaces (Figure 6A). Here, ErbB2 was arranged at the plasma membrane following the micropattern on the surface. When the receptor is activated, downstream signaling adaptor proteins (Grb2 or Shc1) are recruited by ErbB2 kinase activity to receptor-enriched micropatterns (Figure 6B). To quantify the receptor activation state, we used the respective fluorescence signal intensities within and outside the antibody-patterned regions: the degree of ErbB2 activation is expected to be proportional to the level of Grb2/Shc1 corecruitment to the active receptor variants, which is in turn reflected in the normalized mRFP-contrast value (see Supplementary Methods). False-positive TIRF signals due to the deformation of the plasma membrane on top of the patterned antibody surfaces were excluded since control cells cotransfected with Lact-C2-RFP (which was fused with the C2 domain of bovine lactadherin) showed no patterning of the RFP signal due to homogenous membrane distribution in the central regions of the GFP-ErbB2-patterned cells, as shown in detail in (Karimian et al. 2022) and Supplementary Figure S11.

**Figure 6.**
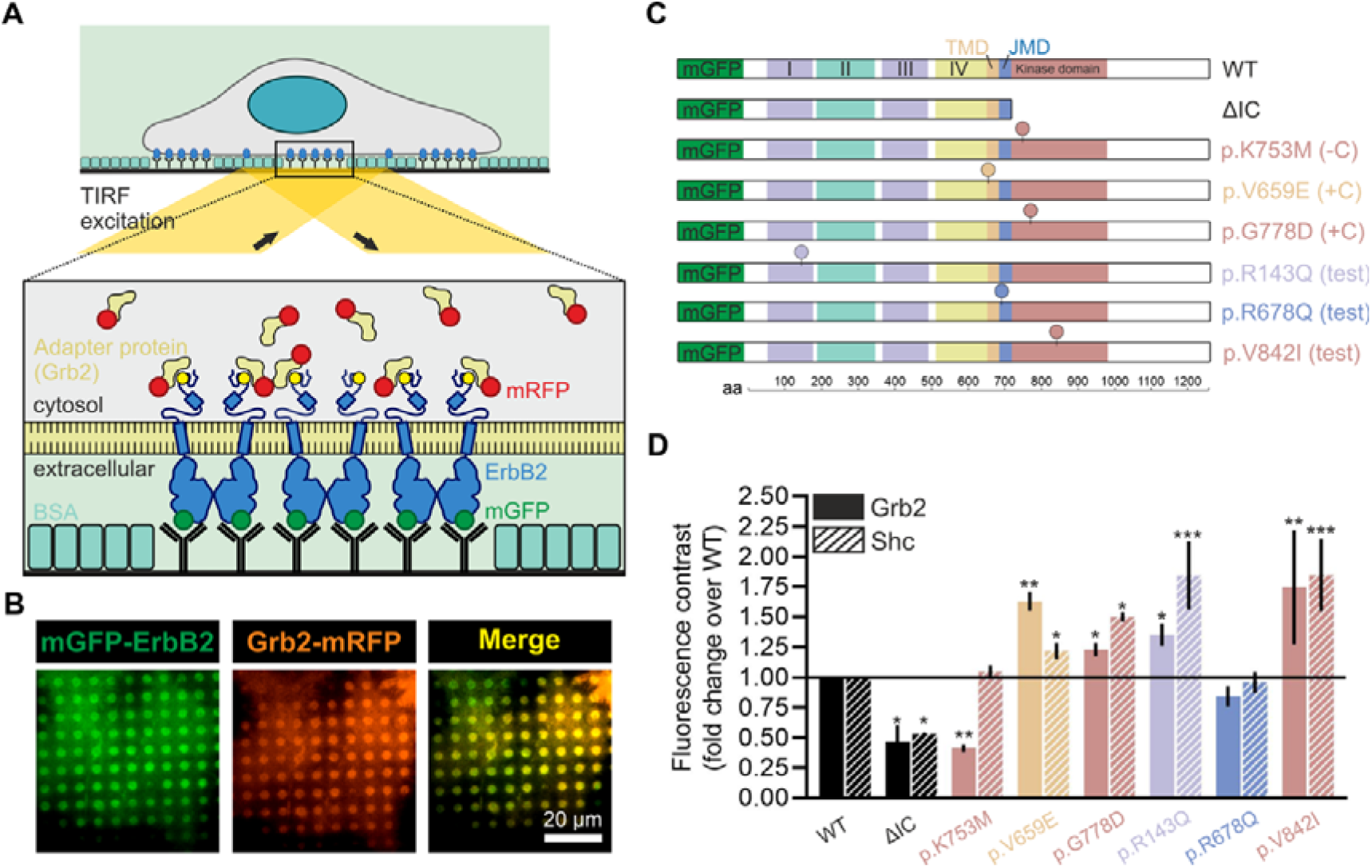
Functional signaling consequences of ErbB2 mutations. **A.** Schematic representation of the micropatterning assay. The cells were transiently transfected with fluorescently labeled bait (mGFP-ErbB2) and prey (Grb2-mRFP, Shc-mRFP) molecules and grown on anti-mGFP antibody-patterned surfaces. The colocalization of the respective adapter protein with mGFP*-*ErbB2-enriched areas indicates the activation state of the receptor. TIRF microscopy is used to specifically detect membrane-proximal fluorescent proteins and to reduce cytosolic background signals. **B.** Representative TIRF images of HeLa cells transiently coexpressing mGFP-ErbB2 (WT) and Grb2-mRFP grown on anti-mGFP antibodies. **C.** Schematic representation of fluorescent ErbB2 fusion proteins and variants used for activation analysis. The positions of the variants are indicated at their approximate locations in the protein domains with their respective amino acid substitutions. **D** Quantitation of bait-normalized fluorescence contrast of Grb2 and Shc adapter protein copatterning. The data were normalized to those of the WT control and are presented as the mean ± SEM (n > 45 cells measured on at least three different days). All measurements were carried out in the nonliganded state, as there is no active or specific ligand known for ErbB2. Only significant differences between the WT and the respective variants are indicated. The *P* values are represented as follows: p≤0.05 (*), p≤0.01 (**), and p≤0.001 (***).

With this approach, we measured the activation of our focal ErbB2 variants and compared them to that of wild-type ErbB2; two negative controls [-C; a kinase dead variant (p.K753M) (Klos et al. 2006); a truncated mutant lacking the intercellular domain (ΔIC); and two positive controls (+C) [constitutively active variants p.V659E and p.G778D (Tan et al. 2005; Klos et al. 2006; Fan et al. 2013; Lorch et al. 2019)]. The signaling activity normalized to that of the wild type is shown for all eight analyzed variants for both Grb2 and Shc1 adapter protein corecruitment (Figure 6C-D, Supplementary Figure S10 and Supplementary Table S9 and S10). None of the negative controls corecruited Grb2. The negative control results for Shc1 were mixed: ΔIC had a reduced interaction with Shc1, but that of p.K753M(-C) remained unchanged, consistent with Shc1 having a different binding site, as also reported by (Schulze et al. 2005). Both positive controls (p.V659E and p.G778D) presented ∼1.6-fold and ∼1.2-fold increased recruitment of Grb2 and ∼1.2-fold and ∼1.5-fold increased recruitment of Shc1, respectively. Of the three focal variants, p.R678Q, the reoccurring variant in cancer patients, did not show increased activity, while both p.R143Q and p.V842I reported significantly greater receptor activity than the wild-type, ∼1.4-1.8-fold, similar to that of the constitutively active positive controls.

## Discussion

Approximately 80% of the new germline mutations passed through generations can be traced back to the paternal lineage (Kong et al. 2012; Francioli et al. 2015). Therefore, investigating mutations that occur in the male germline during a lifetime can yield valuable insights into human diseases, their potential impact on future generations, and evolutionary processes (Campbell and Eichler 2013; Rahbari et al. 2016; Moore et al. 2021). Identifying these mutations has been challenging due to their low frequency.

Using ecSeq, our study examined the occurrence of *ErbB2* DNMs in the male germline, which resulted in an extensive dataset of mutations enriched in sperm DNA from donors of varying ages. This approach facilitated the characterization of the mutational distribution, spectra and signatures specific to *ErbB2*, thereby furthering our understanding of driver mutations and their expansion in the male germline. In addition, this large dataset had sufficient power for testing the preferential accumulation of nonsynonymous versus synonymous substitutions (*d*_N_/*d*_S_ analysis), which identified positive selection to explain the enrichment of missense *ErbB2* mutations observed in sperm DNA. The data also revealed that mutations accumulate even at a young age and are likely to exist as micromosaics within mutation pockets that remain relatively small and constant over time, with no large expansions measured in the aged testis, a novel finding for driver mutations. Functional analyses using biophysical methods further demonstrated that selected ErbB2 mutations hyperactivate the RTK pathway, leading to increased downstream signaling, which likely contributes to the accumulation of ErbB2 mutations in the male germline. Our findings provide relevant insights into the enrichment of mutations or micromosaicism in the male germline, impacting the transmission and recurrence of ErbB2-associated disorders independently of age.

### Origin of *ErbB2* mutations

Germline DNMs can arise at various stages, including early embryogenesis, primordial germ cell differentiation, prepubertal or postpubertal spermatogenesis and adulthood, ultimately populating the sexually mature testis (reviewed previously (Rahbari et al. 2016; Goldmann et al. 2019)). Some mutations may be driven by selection, while others may change through the stochastic process of genetic drift (McFarland et al. 2014; Martincorena et al. 2018), resulting in different proportions of mosaicism, as described in hematopoiesis (Acuna-Hidalgo et al. 2016; Acuna-Hidalgo et al. 2017). DNMs linked to selection are enriched by one lineage of cells harboring the mutation being favored over another, producing more daughter cells (Sottoriva et al. 2015; Turajlic et al. 2019). This results in the expansion of DNMs with age, with strong driver mutations leading to selective sweeps in the male germ line. However, the loss or fixation of neutral or passenger DNMs also reduces the number of cell lineages and the variation or heterogeneity of subpopulations, as was observed for white blood cell DNA with age (Salk et al. 2018). Consequently, neutral random drift might be misinterpreted as a selective sweep (McFarland et al. 2014).

The chance of encountering any particular DNM at a given specific site is very low, with an average frequency of ∼10^-8^ in the human genome. The exonic ErbB2 DNMs had a VAF 4-5 orders of magnitude greater than that of the other DNMs, with individual donors showing frequencies as high as ∼5×10^-4^. How can such a high VAF be explained? Furthermore, how does this explanation fit the observation that 10% of the exonic *ErbB2* DNMs occurred independently in multiple donors, particularly missense mutations occurring at both CpG and non-CpG sites? An explanation based solely on a “hypermutable site” concept (with CpG transitions having only one order of magnitude higher frequencies than non-CpG transitions) is unsatisfactory. Random drift cannot explain these high frequencies either. The probability of encountering the same high-frequency mutation occurring independently in different donors by random chance or drift is exceedingly small. A more plausible scenario is that these mutations arise infrequently but cause a functional change coupled with a growth advantage and undergo clonal expansion within the male gonad, leading to a relative enrichment of mutant spermatogonia or their sperm equivalents, characteristic of driver mutations.

Furthermore, our large dataset had sufficient power for testing the preferential accumulation of nonsynonymous versus synonymous substitutions (d_N_/d_S_ analysis) and identifying positive selection as an explanation for the enrichment of missense ErbB2 mutations in sperm DNA. This trend was more robust when considering recurrent mutations. Notably, recurrent mutations were mainly C>T substitutions at CpG sites, in contrast to the complete dataset, which was enriched for non-CpG transitions and transversions. A similar difference was observed between recurrent mutations (sibling-shared mutations) derived from the maternal lineage or paternal lineage (Jonsson et al. 2018), the latter being more similar to recurrent *ErbB2* mutations.

The observed recurrent substitutions were mainly missense substitutions in the kinase domain and were described as deleterious by different predictors (CADD, SIFT and PolyPhen scores). These mutations might be linked to changes in signaling activity, as was shown for selected RTK signaling pathway activation by receptor-adaptor interactions at the cell membrane of live cells (micropatterns combined with TIRF) for two of the selected mutants (p.R143Q, p.V842I) that exhibited elevated recruitment of Shc and Grb2 compared to WT ErbB2. Intriguingly, the downstream signaling pathway was similar to that of the WT R678Q mutant. These results are in agreement with previous findings (Bose et al. 2013). In light of these results, we hypothesize that the *ErbB2* variants captured with ecSeq in sperm DNA are enriched in the male germline by positive selection.

#### Are ErbB2 mutations exclusive to the sexually mature germ line?

The ongoing cell divisions at sexually mature male gonads might lead to the accumulation of driver mutations with age. For decades, it has been hypothesized that the expansion of driver mutations occurs exclusively in the sexually mature male germline (adulthood), explaining the larger mutation pockets in aged testes but not in the gametes of young donors (Choi et al. 2008; Choi et al. 2012; Yoon et al. 2013; Arnheim and Calabrese 2016). The mutation rarely arises but expands clonally in the sexually mature testis, leading to a relative enrichment of mutant spermatogonia or sperm equivalents. Critical for the formation of these clusters is that all of the descendants stay in close proximity to the initial mutant cell (reviewed in (Arnheim and Calabrese 2009; Arnheim and Calabrese 2016)).

However, the expansion of driver mutations might not be limited to postpubertal spermatogenesis and adulthood but might occur at earlier stages, including early embryogenesis, primordial germ cell differentiation, or prepubertal spermatogenesis. Recent work also revealed increased FGFR3 VAF frequencies in young sperm donors (Salazar et al. 2022; Moura et al. 2024). In particular, missense mutations were observed in 0.01-0.005% of FGFR3 patients, and in some cases, the frequency did not increase with donor age (Moura et al. 2024). In particular, for *FGFR3,* a well-studied driver gene, the studied gain-of-function variants with promiscuous activation (ligand-independent) exhibited two distinct mutational behaviors: one that grows to larger subclonal clusters in the sexually mature gonad and one that increases in frequency in sperm with age. The other likely occurs before puberty, during which stable niches that stay constant in size are formed (as also described here for ErbB2), challenging the long-standing hypotheses that driver or selfish mutations originate exclusively in the sexually mature male germline and continue growing with time (Moura et al. 2024).

The accumulation of variants before postpubertal spermatogenesis was also reported in family pedigrees for neutral mutations, with DNMs being shared among siblings and coming from the same parent (Rahbari et al. 2016) but not present in the parent’s somatic cells and with no evidence for a dependency on parental age (Gao et al. 2019). Additionally, for other species, including reptiles, birds, and mammals, mutations reportedly accumulate not only during spermatogenic cycles postpuberty but also during earlier developmental phases (de Manuel et al. 2022).

The hypothesis that mutations can accumulate in the germline before puberty would also align with the expansion patterns observed for *ErbB2*. We found evidence for positive selection (d_N_/d_S_ analysis) in all three age categories (younger, middle and older donor groups), mainly with variants in the extracellular and protein kinase domains, indicating likely functional consequences.

Our testicular dissection study also suggested that some *ErbB2* mutations establish stable niches that remain constant in size for the three selected ErbB2 variants (c.428G>A, c.2033G>A, and c.2524G>A). This stable size may be attributed to the fact that some variants are tolerated only at low levels, as often observed in highly activating rasopathies in which mosaics are formed in the skin (Tiemann-Boege et al. 2021). A similar behavior was reported for selected *FGFR3*-activating mutations, which also showed increased frequencies in young donors that formed rather small mutation pockets in the testis (Moura et al. 2024) and for early developmental neutral clones that remained temporally stable across serial samples and age groups, with no changes in size (or frequency) in the stem cell niches (Yang et al. 2021). This also challenges the dogma that driver mutations expand into large clusters with time in the male germ line.

In conclusion, while age-associated driver mutations are more prevalent in offspring from fathers of advanced age, the risk of recurrence in siblings or the incidence in the general population might be greater for germline mosaics than for age-associated mutations; however, this might strongly depend on the selective advantage conferred by the mutation. This is particularly worrisome since different ErbB2 activating mutations might have early- or late-onset effects associated with a clinical phenotype that ranges from a rare genetic disorder to cancer or tumor resistance to certain protein tyrosine kinase inhibitors.

## Materials and Methods

### Sample collection, preparation and DNA extraction

Sperm samples from anonymous donors who were abstinent for > 3 days were collected at Kinderwunsch Klinik, MedCampus IV, Kepler Universitätsklinikum, Linz following the protocol approved by the ethics commission of Upper Austria (Approval F1-11). The donors were mainly between 19 and 63 years old and were mostly of European ancestry. Two snap-frozen, postmortem testes from 73-year-old (ID: NRD#ND 10354) and 70-year-old (ID: NRD#ND 10225) donors were collected from the National Disease Research Interchange (NDRI, Philadelphia, PA). None of the donors had chronic infections; diabetes; chemotherapy; radiation; or alcohol, tobacco, or drug abuse. DNA was extracted from fresh semen samples following the protocol described previously (Arbeithuber et al. 2015; Moura et al. 2024). Testis dissection was performed as previously described (Qin et al. 2007; Choi et al. 2008; Choi et al. 2012; Shinde et al. 2013; Yoon et al. 2013; Moura et al. 2024; Striedner et al. 2024). The details of the sperm and testis DNA extraction can be found in the SM Methods. Information about the different sperm donors, including the WHO classification for human semen characteristics (Cooper et al. 2010) and the library protocol used, is listed in Supplementary Table S11.

#### Library preparation

Library preparation was performed according to the protocol of (Salazar et al. 2022), with modifications from (Loeb et al. 2019; Abascal et al. 2021), as outlined in Supplementary Table S12 and the Supplementary Materials. We used two strategies: repair of the libraries involved treatment with the USER enzyme (NEB, M5505S) and Fpg-glycosylase (M0240S, NEB), along with a blunting step using mung bean nuclease (NEB, M0250S) and ddBTPs (Merck/Sigma Aldrich, GE27-2045-01) for nick sealing, as detailed in the Supplementary Methods and Supplementary Table S12. For unrepaired libraries, we used focused ultrasonication (Covaris M220 instrument) followed by size selection, end repair, and A-tailing.

For adapter ligation, DS_Hairpin_U adaptors with 12 random nucleotides were used for ligation via the NEBNext Ultra II end repair/dA-tailing module (NEB) and the NEBNext Ultra II ligation module for both library types. The USER enzyme digest (NEB) was used to open the adapter loop. Adapters were synthesized as previously described (Salazar et al. 2022) and are specified in the Supplementary Methods. Amplification was performed using variable input DNA, and the PCR conditions involved 12 or 6 cycles of single primer extension followed by 2 PCR cycles. The reaction was carried out in 1x Kapa HiFi Reaction Mix (Roche) followed by DNA cleanup with 1.2x volumes of Sera-Mag Select beads (Cytiva) according to the manufacturer’s instructions. The input DNA, PCR conditions, and reaction volumes are described in the Supplemental Methods and Supplementary Table S13. The primer sequences and oligonucleotides used are shown in Supplementary Table S14. In both strategies, after initial amplification, 120 bp biotinylated oligonucleotide probes were employed for two rounds of targeted capture followed by further PCR cycles for sufficient target enrichment, as described previously (Schmitt et al. 2015; Salazar et al. 2022). Details, including the number of cycles for each capture PCR and the sequence of the biotinylated oligonucleotide probes, are provided in Supplementary Tables S13, S14, S15, and S16 and in the Supplemental methods. Sequencing was performed using the MiSeq Reagent v3 600 cycles kit at the VBCF NGS Unit, Vienna, Austria, or primarily on the HiSeq X 150 PE platform at Macrogen, Seoul, South Korea.

#### Duplex Sequencing Data Analysis and Variant Filtering

The raw sequencing data were analyzed with the Du Novo package on the Galaxy platform (https://usegalaxy.org/u/jku-itb-lab/w/galaxy-workflow-ErbB2, Supplementary Figure S12, a duplex sequencing (DS) pipeline that assembles reads sharing the same barcode into families before alignment to the human genome assembly hg38 (Stoler et al. 2016). For details on the pipeline, see the SM Methods. We also removed the first and last 15 nucleotides of the duplex consensus sequence (DCS) as potential artifacts deriving from error-prone end repair. Additionally, we used the Variant Analyzer tool (VAR-A) (Povysil et al. 2021) to classify the confidence in the identified alternate alleles by a tier-based system, as described in Supplementary Table S17. We filtered all intronic variants and SNPs from the analysis, and only variants in high-quality tiers (tiers 1.1-2.5) were retained. We also removed variants that cooccurred with another rare variant or rare haplotype and within STRs. This list of variants is denoted as “all variants”. Moreover, “recurrent variants” represent variants that occur in at least two libraries.

Finally, the variants were annotated with wANNOVAR (Yang and Wang 2015) and the variant effector predictor (VEP) (McLaren et al. 2016). The deleteriousness of a variant was described with the CADD (Rentzsch et al. 2019) and SIFT scores. Additionally, associations with a cancer type were extracted from COSMIC, and *ErbB2* variants (human genome assembly hg38) of the transcript ENST00000269571.5 were extracted from gnomAD v3.1.2 (https://gnomad.broadinstitute.org) and Cosmic V96 (https://cancer.sanger.ac.uk).

#### Substitution frequency

The substitution frequency was calculated as the number of de novo events per number of sequenced nucleotides, which was estimated as the mean coverage multiplied by the targeted region (exonic size). If a mutation occurred in different libraries, it was also counted multiple times.

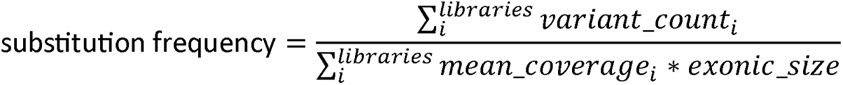

#### Mutational spectra

We categorized the mutational spectra after the (un)transcribed strand. The mutation spectra are estimated with relative unique counts, where each variant is counted only once, regardless of the number of occurrences in the libraries, to compare it to the variant counts from public databases. Only substitutions of the *ErbB2* gene (transcript ENST00000269571.9) and within the negative control regions (human genome assembly hg38) were extracted from the gnomAD v3.1.2 (Karczewski et al. 2021) and COSMIC v96 (Tate et al. 2019) databases.

#### Cosine similarity

The similarity between a query and reference mutational spectra is measured with the cosine similarity. First, a distribution of cosine similarities is generated by randomly sampling n mutations (number of mutations of the query spectra) from the reference spectra and each of the samples is compared to the original reference spectra (1,000 iterations). Second, the observed cosine similarity can be compared to the 95% confidence intervals of the bootstrapped reference samples. This approach is followed as in (Abascal et al. 2021).

#### Mutational signature

The mutational signatures are built from the trinucleotide context (5’ and 3’ neighboring nucleotide of the variant) with the tools SigProfilerMatrixGenerator and SigProfilerPlotting (Bergstrom et al. 2019)and compared to the COSMIC signatures v3.3 (Alexandrov et al. 2020) with SigProfilerExtractor (Salazar et al. 2022).

#### The ratio of nonsynonymous to synonymous variants

The *d*N/*d*S ratio was estimated as described previously (Martincorena et al. 2017) using the dNdScv package v. 0.1.0 in R v. 4.3.1 and the human reference sequence v. GRChg38.14 and annotation for the ErbB2 region v.GR38.111 from ENSEMBL (https://www.ensembl.org/). In this analysis, the dN/dS estimates considered the nearby sequence context but not the genome-wide epigenetic covariates of the mutation rate (Nei and Gojobori 1986).

#### Droplet Digital PCR (ddPCR)

The fractional abundance (FA) of mutations was evaluated by ddPCR (Bio-Rad) (Hindson et al. 2011) as described previously (Moura et al. 2024). The site-specific mutation detection assays were designed using the online platform of Bio-Rad (https://www.Bio-Rad.com/digital-assays) listed in Supplementary Table S18. For each ddPCR, 10 µl of 10x SuperMix for Probes (no dUTP), 6.7 µl of nucleic acid-free water, 2 µl of genomic DNA (∼125 ng/µl; ∼36,000 genomes/µl), 1 µl of probes (900 M of each probe and 100 M of each primer), and 0.3 µl of restriction enzyme (MseI; 10 U/µl) were added, and the mixture was incubated at room temperature for 15 minutes to enhance target template availability. The digested mixture, along with 70 µl of droplet generation oil, was loaded into a cartridge covered with a gasket to then form the droplets in the droplet generator (Bio-Rad). Approximately 43 µl of the droplet solution was transferred to a ddPCR 96-well plate and sealed in a PX1 PCR plate sealer (Bio-Rad) followed by PCR at 95°C for 10 minutes; 40 cycles of 94°C for 30 seconds and 53-56°C (see Supplementary Table S18) for 1 minute; and a final step of 98°C for 10 minutes, with a ramp rate of 2°C/sec and a lid temperature of 105°C. The end-point PCR plate was subjected to data analysis using QuantaSoft Analysis Pro Software (version 1.7.4; Bio-Rad Laboratories, Inc.). The threshold between positive and negative droplets was manually adjusted well by well or across an entire plate based on the fluorescence amplitude (fluorescence intensity of positive and negative droplet clusters and/or histogram plots) for each specific probe, and we reported here the Poisson-corrected data points.

### Site-Direct Mutagenesis (SDM)

To assess the functional impact of the selected mutations, site-specific single-nucleotide mutations were introduced into the mGFP-ErbB2 expression plasmid (Szabo et al. 2022) using a site-directed mutagenesis strategy with Phusion HS II high-fidelity polymerase as detailed in the Supplementary Methods and Supplementary Table S19 and described previously (Hartl et al. 2023; Moura et al. 2024). The designed oligonucleotide pairs, including a primer with the desired mutation and another primer with a silent mutation, both featuring 3’ PTO bonds for enhanced specificity, were used for amplification. The resulting PCR products, which were confirmed to be of the correct length via gel electrophoresis, were ligated without purification, followed by DpnI digestion and transformation into *E. coli* NEB 10-beta. Ampicillin-resistant clones were screened by colony PCR, and positive clones were cultured overnight, subjected to plasmid miniprep, and sequenced to verify the introduced mutations.

### Live*-*cell micropatterning experiments

Preparation of protein micropatterned surfaces by large-area microcontact printing (Karimian et al. 2022), total internal reflection fluorescence microscopy (Lanzerstorfer et al. 2014), and image analysis (Hager et al. 2021) was carried out as previously reported and is described in the Supplementary Methods and Supplementary Figure S11.

#### Data Access

The raw sequencing data generated in this study have been submitted to the NCBI BioProject database (https://www.ncbi.nlm.nih.gov/bioproject/) under accession number PRJNA1052412.

## Acknowledgments

We would like to thank Václav Brož for his help in the testis DNA extraction and measurements with ddPCR and Philipp Hermann for advice on the statistical testing.

## Authors’ contributions

T.M., A.Y., T.K., and I.H. performed the experiments. M.H., S.M., P.L., A.J.B. and I.T.-B. analyzed the data. I.T.-B. conceived the project and provided funding. A.Y., M.H., T.M., T.K., P.L., A.J.B., and I.T.-B. wrote the manuscript. All the authors read and approved the final manuscript.

